# Causal contribution and dynamical encoding in the striatum during evidence accumulation

**DOI:** 10.1101/245316

**Authors:** Michael M. Yartsev, Timothy D. Hanks, Alice M. Yoon, Carlos D. Brody

## Abstract

A broad range of decision-making processes involve gradual accumulation of evidence over time, but the neural circuits responsible for this computation are not yet established. Recent data indicates that cortical regions prominently associated with accumulating evidence, such as posterior parietal cortex and the frontal orienting fields, are not necessary for computing it. Which, then, are the regions responsible? Regions directly involved in evidence accumulation should satisfy the criteria of being necessary for accumulation-based decision-making behavior, having a graded neural encoding of accumulated evidence and causal contributing throughout the accumulation process. Here, we investigated the role of the anterior dorsal striatum (ADS) in a rodent auditory evidence accumulation task using a combination of behavioral, pharmacological, optogenetic, electrophysiological and computational approaches. We find that the ADS is the first brain region known to satisfy these criteria. Thus, the ADS may be the first identified node in the network responsible for evidence accumulation.

## Introduction

All behaving animals must interpret sensory information arriving from the environment and use that information to select future actions. How the nervous system solves this problem has been a long-standing question in neuroscience. Multiple studies across a wide range of behavioral tasks and model systems, including humans (Hunt et al., 2012; Krajbich et al., 2012; Ratcliff and Smith, 2015), non-human primates (Gold and Shadlen, 2007; Huk and Shadlen, 2005; Shadlen and Newsome, 1996) and rodents (Brunton et al., 2013; Carandini and Churchland, 2013; Erlich et al., 2015; Hanks et al., 2015; Raposo et al., 2012) have proposed a framework by which neural circuits gradually accumulate sensory evidence to guide decisions. Yet, despite the observation of neural correlates of evidence accumulation in several brain regions (Ding and Gold, 2010; Gold and Shadlen, 2007; Hanks et al., 2015; Ratcliff et al., 2007; Shadlen and Newsome, 1996), a major challenge of this line of research has been that the neural circuits causally responsible for evidence accumulation have not yet been determined. Studies of two of the cortical regions most prominently associated with evidence accumulation, posterior parietal cortex (PPC; Huk and Shadlen, 2005; Kira et al., 2015; Roitman and Shadlen, 2002; Shadlen and Newsome, 1996) and the frontal eye fields in primates (FEF; Ding and Gold, 2012a; Gold and Shadlen, 2000; Mante et al., 2013) together with its likely rodent analogue, the frontal orienting fields (FOF; Erlich et al., 2011) have recently presented evidence indicating that neither the PPC nor the FOF is central to the computation of gradually accumulating evidence (Erlich et al., 2015; Hanks et al., 2015; Katz et al., 2016). The anterior dorsal striatum serves as an intriguing alternative candidate, due to its unique anatomical positioning as a convergence hub for multiple brain regions (such as the PPC and FOF) where neural signatures of evidence accumulation have been observed (Cheatwood et al., 2003; Ding and Gold, 2013a; McGeorge and Faull, 1989). It is thus ideally positioned to participate in evidence accumulation as part of its established role in action selection (Bogacz and Gurney, 2007; Graybiel, 2008; Hikosaka et al., 2014; Jin and Costa, 2010; Nelson and Kreitzer, 2014; Redgrave et al., 2010). The auditory input to a different striatal subregion, the posterior “auditory” striatum, has been shown to be critical for auditory discriminations, leading to the suggestion that cortical projections into the striatum may provide a general mechanism for the control of motor decisions (Xiong et al., 2015; Znamenskiy and Zador, 2013). Moreover, recent experimental work has suggested that the anterior dorsal striatum may contribute to the computations specifically involved in evidence accumulation (Ding and Gold, 2010, 2012b, 2013b; Lo and Wang, 2006). Yet three critical tests of this idea have been left unanswered. First, is the dorsal striatum necessary for accumulation-based decision-making? To date, there have been no inactivations of the dorsal striatum during accumulation of evidence. Inactivations are critical tests of whether a region reflects a variable, or can be determined to play a causal role in computing it (Erlich et al., 2015; Katz et al., 2016). Second, do neurons in the dorsal striatum encode sensory information in a sufficient way to be directly involved in the graded accumulation process? The correlates of evidence accumulation reported so far in striatum are firing rates that, when averaged over trials, ramp upwards, where the slope of the ramp grows with increasing strengths of the evidence (Ding and Gold, 2010, 2012b). But these trial-averages are also consistent with other encodings that on a single-trial basis do not represent gradually accumulating evidence, such as sharp steps in firing for which the timing of the step varies across trials (Hanks et al., 2015; Latimer et al., 2015). Thus we have only indirect information as to whether the anterior dorsal striatum does or does not encode gradually accumulating evidence. Lastly, does the dorsal striatum play a causal role throughout the period of accumulation? If the striatum is part of the accumulation process that drives behavior, perturbing it at any period during the accumulation process should affect behavior. This feature is thus an essential prerequisite for a component of the accumulator. Yet no temporally-specific perturbations of the dorsal striatum during accumulation of evidence have yet been carried out. No brain region studied during an accumulation of evidence behavior has been reported to possess this feature. Here, using a combination of behavioral, pharmacological, optogenetic, electrophysiological and computational approaches, we address these three fundamental questions and provide evidence supporting a central role for the anterior dorsal striatum in evidence accumulation.

## Results

We trained rats on a previously developed decision-making task (Brunton et al., 2013) in which subjects accumulate auditory evidence over many hundreds of milliseconds to inform a binary left-right choice (Fig. 1a). On each trial, rats kept their nose in a central port during the presentation of two simultaneous trains of randomly-timed auditory clicks, one played from a speaker to their left and the other from a speaker to their right. At the end of the auditory stimulus, the rat’s task was to decide which side had played the greater total number of clicks. Consistent with previous studies using this task, analysis of our rats’ behavior indicated that they gradually accumulated auditory evidence over the entire trial, and used that accumulated evidence to drive a categorical choice (Extended Data Fig. 1; Extended Data Table 1).

**Fig. 1.**
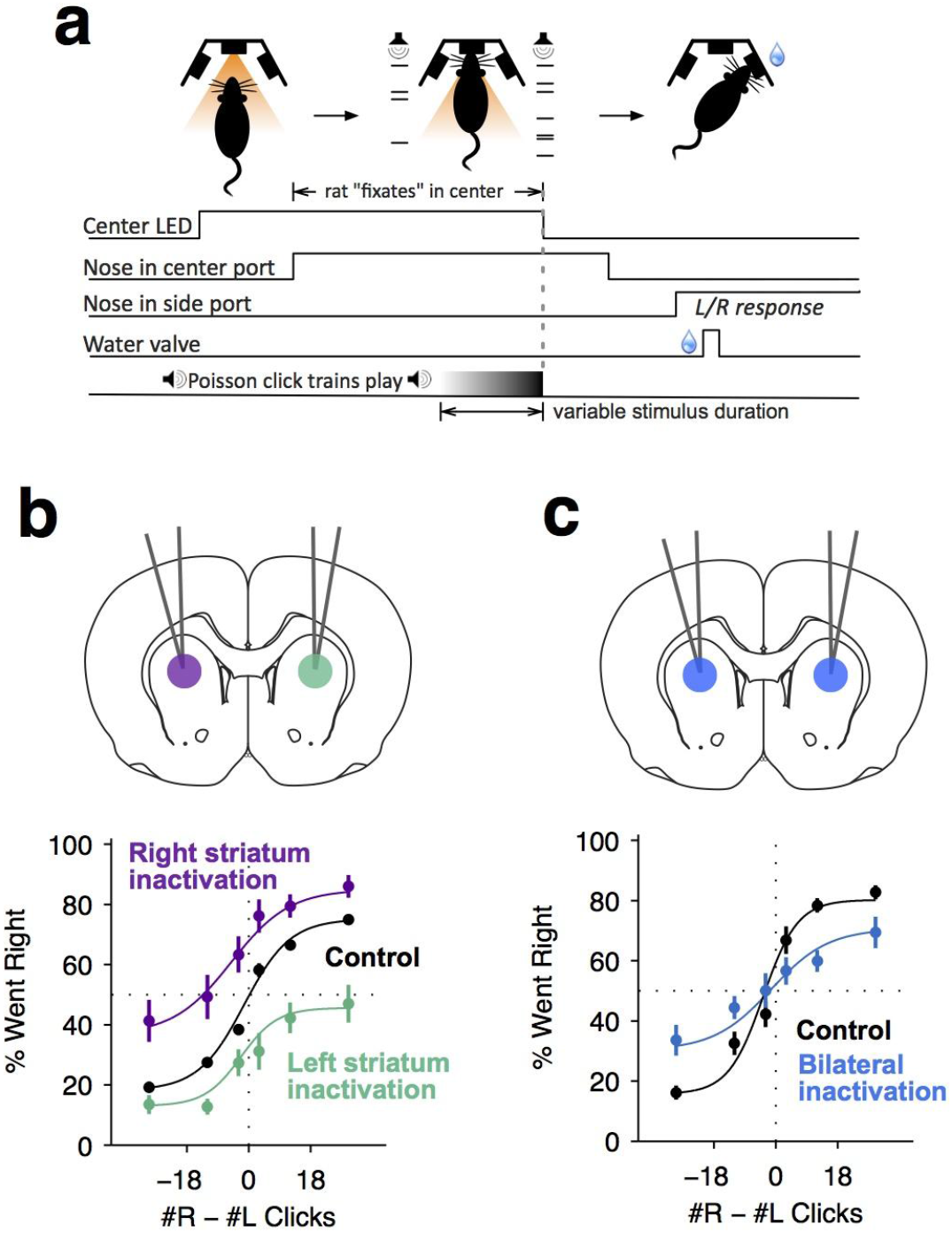
Dorsal anterior striatum is necessary for performance on the Poisson-clicks evidence accumulation task. **(a)** Sequence of events in each trial of the rat auditory Poisson-clicks task. From left to right: after light onset above the center port, rats “fixate” their position by placing their nose inside the center port. During nose-fixation, two different trains of randomly timed auditory clicks are played concurrently from the left and right speakers. Upon termination of the sound trains, the light above the center port turns off and the rat needs to make a choice, poking into the left or right port to indicate if more clicks were played on the left or right sides, respectively. **(b)** Unilateral infusion of muscimol into the striatum results in a significant ipsilateral bias on accumulation trials. Purple and cyan psychometric curves show data on days of right and left striatal infusions (n left sessions = 29; n right sessions = 29), respectively. Black psychometric curve shows data from control sessions that occurred one day before infusion sessions (n = 58). **(c)** Bilateral infusion of muscimol into the striatum results in significant impairment on accumulation trials. Blue psychometric curve is from bilateral infusion sessions (n = 26) and black psychometric curve is from control sessions that occurred one day before bilateral infusion sessions (n = 26). Data is shown as mean ± S.E.M.

We began assessing the role of the anterior striatum in the accumulation task using reversible pharmacological inactivation with muscimol (Methods). The anterior striatial region targeted in this study receives convergent inputs from the PPC and the FOF, brain regions previously reported to contain neural correlates of evidence accumulation but later shown to not be central to the accumulation process itself (Erlich et al., 2015; Hanks et al., 2015; Katz et al., 2016). Unilateral inactivation biased rats to make more ipsilateral choices relative to controls (Fig. 1b; bias for right side inactivation = 19.2 +− 4.4%, p < 0.01; bias for left side inactivation = 18.6 +− 3.3%, p < 0.01). This effect was not a gross motor bias, but was instead specific for accumulation trials, for no significant bias was caused on interleaved motor control trials in which the rats had to make similar left-right motor response, but were cued by a simple visual stimulus (Extended Data Fig. 2; p > 0.4 for both left and right side trials). Bilateral pharmacological inactivation caused a substantial impairment in performance for accumulation trials (Fig. 1c; impairment = 12.6 +− 3.2%, p < 0.01). This impairment was again specific for accumulation trials, with no significant impairment for motor control trials where the decision was not based on the accumulation of evidence over time (Extended Data Fig. 2; p > 0.6 for both left and right side trials).

**Fig. 2.**
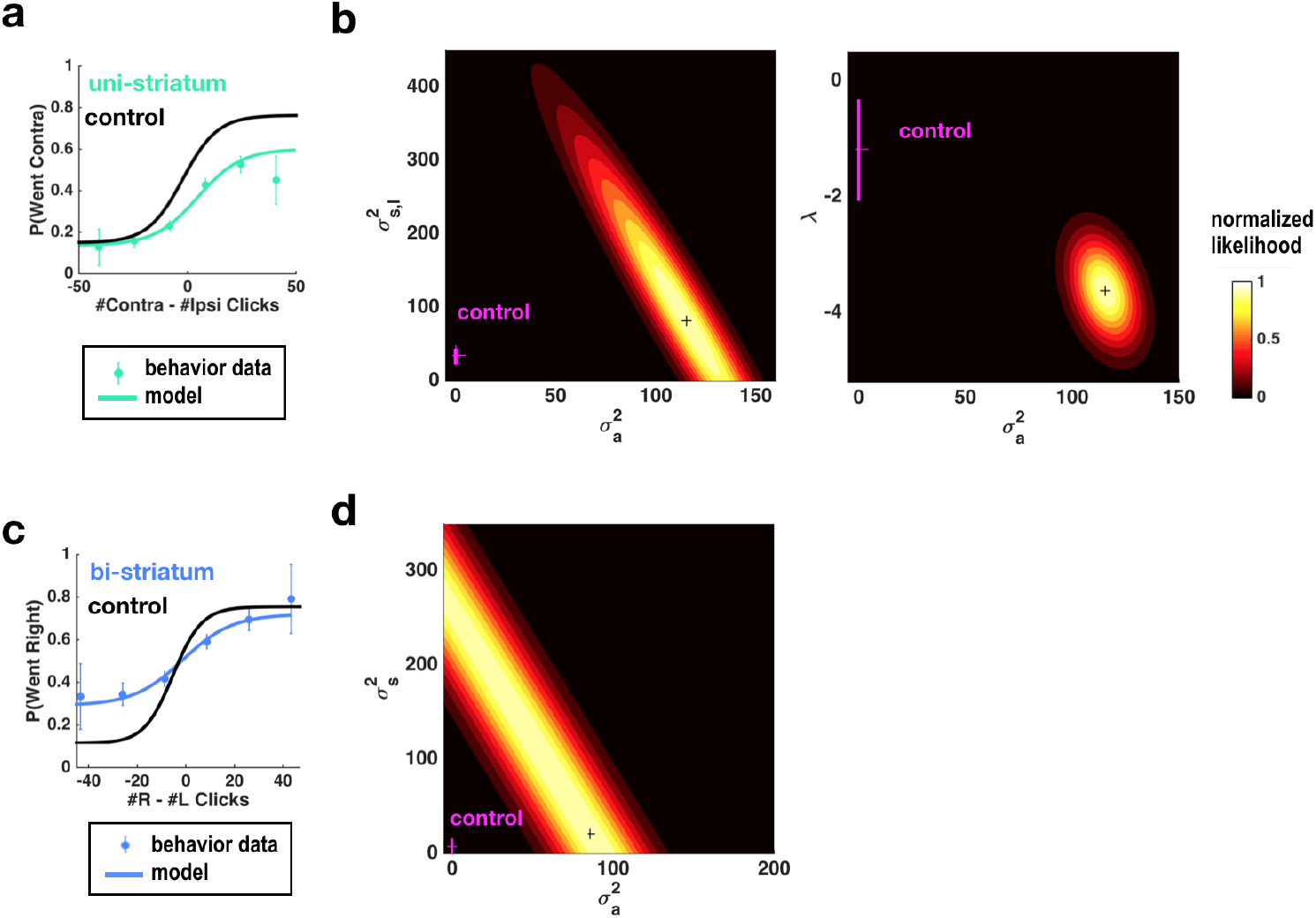
**Fits of the model of** (Brunton et al., 2013;Erlich et al., 2015) **to data from sessions following muscimol inactivation of the striatum**. **a**, Psychometric curves for control and unilateral inactivation data. Left and right inactivations were collapsed together. Cyan data points are from sessions following unilateral infusions of muscimol. The black and cyan lines are the psychometric curves predicted from the model fit to the control and inactivation data, respectively. **b**, normalized likelihood of the data given the model as a function of the parameters for which best-fit values for inactivation data were significantly different from best-fit values for control data. Magenta shows the best-fit values for control data, together with error bars. Black cross shows best-fit values for inactivation data Left: Sensory noise for the side contralateral to the infusion versus accumulator noise. Although there is a trade-off between accumulator and sensory noise, the accumulator noise parameter has a best-fit value following unilateral inactivations that is significantly greater than its control best-fit value. Right: leak/instability parameter versus accumulator noise. Behavior has become significantly leakier. **c,d** as in panels **a,b** but for bilateral striatum inactivation data, and in a model where the sensory noise is constrained to be the same for both sides of the brain, so there is only one sensory noise parameter. Here the tradeoff between sensory noise and accumulator noise is large enough that we cannot distinguish whether one or both are significantly different from their control values, but there is nevertheless a significant increase in their sum.

Psychometric curves as shown in Fig. 1b,c group together trials based on the click difference accrued by the end of the stimulus stream and treat all trials within each group as if they were the same. But in our clicks task we have far more information available, since the precise temporal pattern of each individual trial’s click trains is known. We have previously used this information, together with a model that takes into account those known individual click times, to explain our subjects’ behavior in terms of an drift-diffusion process (Ratcliff and McKoon, 2008), enhanced so that we can obtain trial-by-trial, moment-by-moment estimates of accumulating evidence (Brunton et al., 2013). The model converts the incoming stream of each trial of discrete left and right click stimuli into a scalar quantity *a*(*t*) that represents the gradually accumulating difference between the two streams and drives choices: at the end of the stimulus, if *a* is positive (negative), the model prescribes ‘choose right’ (‘choose left’). The rat’s behaviour is used to simultaneously estimate 8 parameters that govern how *a*(*t*) evolves (Methods). These parameters quantify sensory and accumulator noise, leakiness/instability of the accumulation process, a sticky accumulation bound, sensory depression/facilitation, side bias, and lapse rate.

The original model of Brunton et al. was not constructed to explain different types of side biases, so it had only a single parameter able to account for such lateralized effects. By adding three more parameters that could cause different types of side biases, fitting the extended model to behavioral data following a unilateral inactivation, and asking which parameters are most affected relative to control trials, we can estimate which particular aspect of the behavior was impacted by the inactivations. Following this approach in the case of unilateral inactivations of the FOF, we previously concluded that FOF inactivations were consistent with perturbing a process that was not part of evidence accumulation per se, but was instead downstream of the accumulation process and therefore followed it (Erlich et al., 2015; Piet et al., 2017).

Here we improve upon this analysis to apply it to our striatum inactivation data. At the time of the Erlich et al. 2015 study, the complexity of determining the derivative of the model with respect to all 11 of its parameters precluded us from fitting all 11 parameters simultaneously. We instead performed exhaustive scans in the space of two parameters at a time while the other 9 parameters were fixed to their control (no inactivation) values (e.g., Figure 4 in (Erlich et al., 2015)). Since that time, however, algorithmic differentiation packages, which greatly facilitate computing the derivative of arbitrary differentiable models embodied in computer code, have become widely available (Abadi et al., 2016; Baydin et al., 2015; Revels et al., 2016; The Theano Development Team et al., 2016). Using the ForwardDiff package of the language Julia (Revels et al., 2016) to automatically obtain the derivative with respect to all 11 parameters in the model of (Erlich et al., 2015), we have constructed a package that can efficiently and simultaneously fit all 11 parameters in the model. We are publishing this package in open source form, as part of the contribution of the current manuscript (code available at https://github.com/misun6312/PBupsModel.jl). We validated this approach and our previous FOF analysis by fitting all 11 parameters simultaneously to our previous FOF unilateral inactivation data. This new analysis (Extended Data Fig. 3) confirmed the conclusions about the FOF found in (Erlich et al., 2015). Following this conclusion, we next turned to performing the same analysis on the inactivations data collected in the current study for the anterior striatum.

**Fig. 3.**
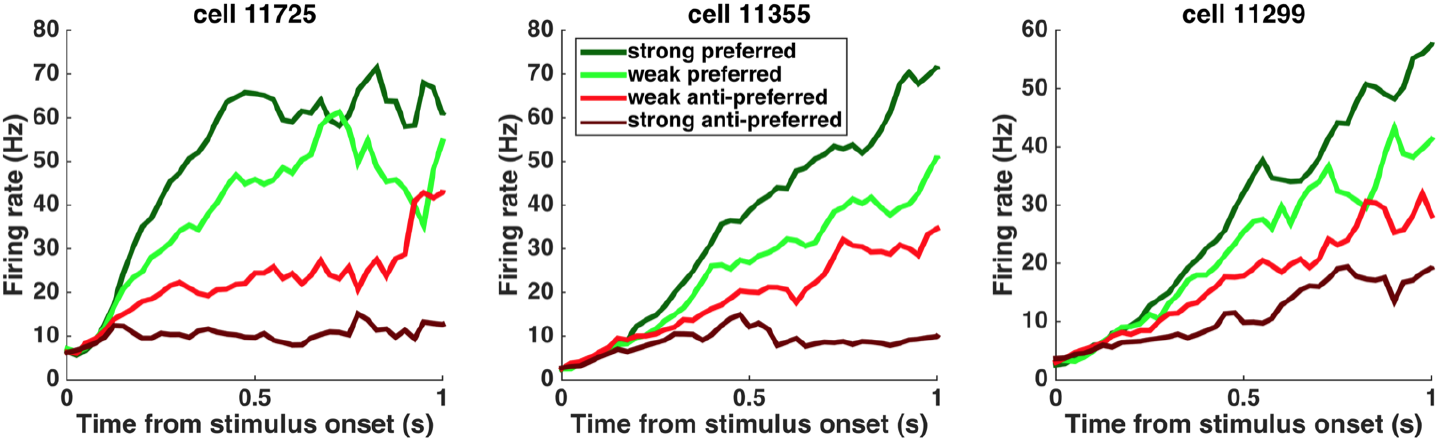
Peri-stimulus time histograms (PSTHs) of example neurons. PSTHs aligned to stimulus onset are shown for 3 example striatum neurons. Trials were sorted into 4 stimulus strength bins for each neuron. Green traces correspond to the preferred-direction stimuli and red traces correspond to anti-preferred-direction stimuli. Darker colors correspond to stronger stimuli (less difficult) and brighter colors correspond to weaker stimuli (more difficult).

**Fig. 4.**
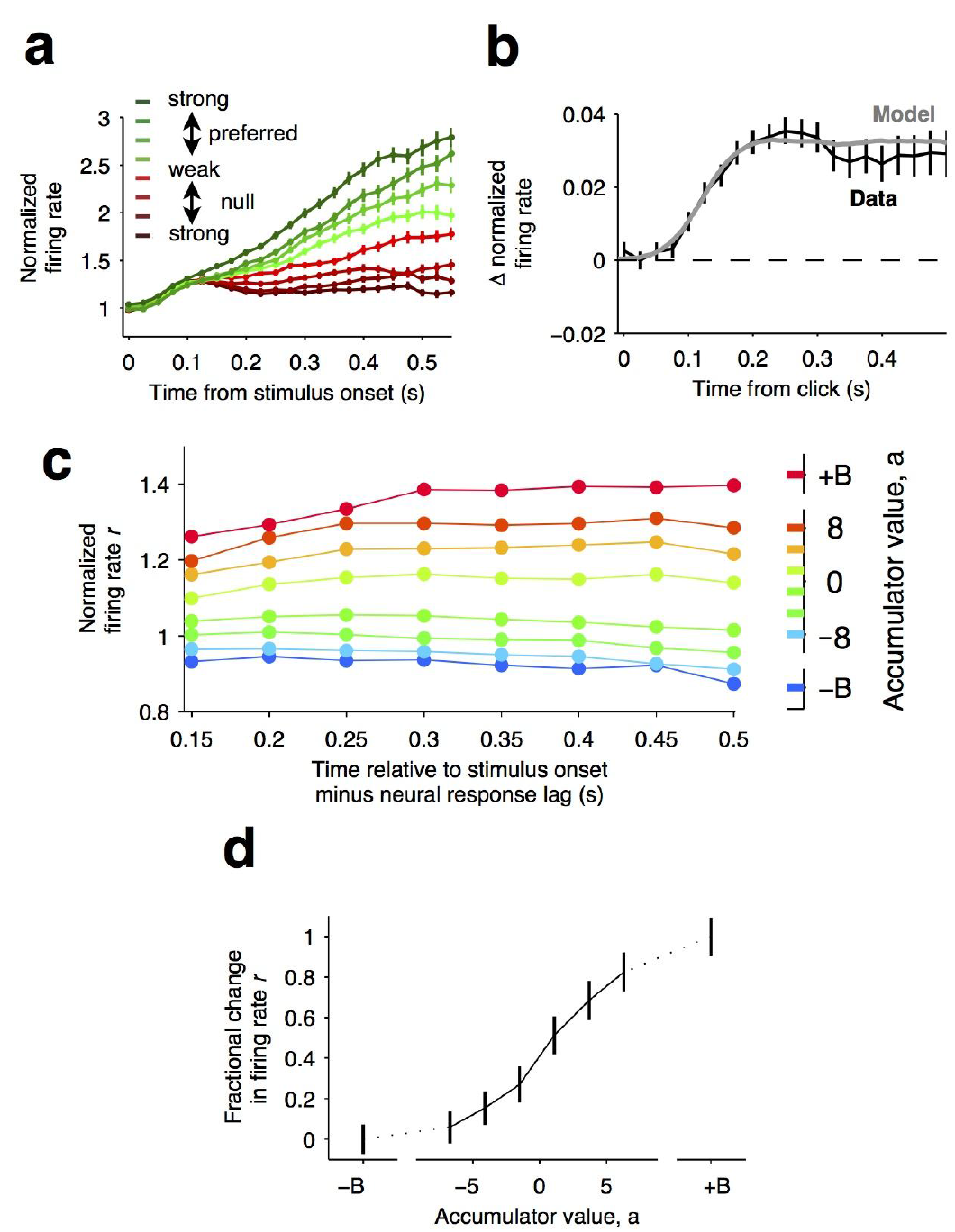
Graded representation of accumulated evidence in the dorsal striatum. **(a)** Responses of pre-movement side-selective striatal neurons during evidence accumulation (mean ± S.E.M.). Trials are grouped by average strength of sensory evidence with greener and redder colors corresponding to stimuli in the preferred and non-preferred direction of the neurons, respectively. Each group of trials is sorted based on difficulty of the trials from easy to hard corresponding to darker and lighter colors, respectively.Note the significant dependence of ramping responses on stimulus strength (n = 64 neurons from 3 rats). **(b)**Click-triggered average response ± S.E.M. Note the close correspondence of the average click-triggered population response to a theoretical prediction of a fixed-magnitude and sustained increase in the neurons’ firing rate (see Methods). **(c)** Firing of striatal neurons aligned to trial onset minus the neural response lag (150 ms; see Methods) grouped based on model-derived accumulator value (colors with ± B correspond to sticky accumulation bounds). Note that this accumulator value to firing rate map is graded and fairly stable over time (n = 64 neurons). **(d)** The population change in firing rate as a function of accumulator value averaged across time exhibits a graded response.

Simultaneously fitting all parameters of the enhanced model to data from sessions with unilateral muscimol inactivation of the anterior dorsal striatum revealed that three parameters differed enough from their control values to produce substantial changes in behavior (Extended Data Table 2). First, the side bias in the lapse rates (difference between the contralateral lapse rate parameter κ_C_ and the ipsilateral lapse rate parameter κ_I_, which are unitless parameters in terms of fraction of trials; Methods) significantly increased in favor of ipsilateral choices (κ_C_ – κ_I_ : from –0.0004 ± 0.020 in control sessions to 0.137 ± 0.0296 for inactivation sessions). An effect on lapse rates was also seen after unilateral FOF inactivations, where it was interpreted as an effect on processes subsequent to the accumulator, and not part of it (Erlich et al., 2015). Second, the accumulator had become leakier (λ : from –1.1795 ± 0.5042 sec ^−1^ in control sessions to –3.6219 ± 0.4463 sec^−1^ for inactivation sessions). Finally, the magnitude of the accumulator noise, which describes diffusion noise intrinsic to the accumulator, also increased significantly (Fig. 2a,b). It was much larger during inactivations than its near-zero value in control sessions (σ^2^_a_ : From 5×10^−6^ ± 0.001 clicks^2^/sec in control sessions to 115.44 ± 38.79 clicks^2^/sec during unilateral inactivations).

For data from bilateral inactivation sessions, the sum of sensory plus accumulator noise was significantly greater than control (Fig. 2c,d; σ^2^_a_ +σ^2^_s_: From 7.37 clicks^2^/sec (95% C.I. = [0 18.16]) in control sessions to 106.72 clicks^2^/sec (95% C.I. = [93.12 361.19]) during inactivations; we note that noise magnitudes cannot be less than zero, implying that confidence intervals for both σ^2^_a_ and σ^2^_s_ are bounded by zero.). But the trade-off between the two (Brunton et al., 2013) was large enough that it was impossible to distinguish whether one or both of σ^2^_a_ and σ^2^_s_ was responsible for the increase. A parsimonious account of the changes in noise parameters for both unilateral and bilateral muscimol inactivations of the anterior dorsal striatum would be that inactivation of the anterior dorsal striatum increases the accumulator noise, but we cannot rule out that the bilateral inactivation data could be due instead to an increase in the sensory noise.

For both unilateral (Extanded Data Table 2) and bilateral (Extended Data Table 3) inactivations, the decision boundary Ϸ, which divides final accumulator values into “go Left” verus “go Right” trials, changed significantly from control values. But the magnitude of this change corresponded to an almost negligible horizontal shift in the psychometric curve, of approximately one click or less (for comparison, the horizontal axis in Fig. 2a ranges from –50 to +50), and for this reason we do not consider Ϸ further here.

These fits contrast with those following FOF inactivation (Erlich et al., 2015). In particular, we note that the sensory and accumulator noise parameters were minimally altered after FOF inactivation, whereas ADS inactivation significantly impacted them.

This pharmacological demonstration that the striatum is necessary for decisions based on the accumulation of evidence, and the model-based suggestion that the striatum affects properties of the accumulator led us to explore the detailed neural dynamics that may support its potential causal contribution. To do so, we conducted single-unit recordings from freely behaving subjects engaged in the evidence accumulation task. Consistent with previous work (Graybiel, 2008; Jin and Costa, 2010; Kravitz and Kreitzer, 2012), we found that the neural activity of many striatal neurons was modulated by movement initiation (Extended Data Fig. 4 A-C). However, we have also found that over a third of the recorded neurons significantly modulate their activity during the fixation period (64/173 of the neurons active during the fixation period; 37%, Extended Data Fig. 4 D-F), many hundreds of milliseconds before the movement initiation reporting the decision. This timing suggests they may have a role in forming the decision, and we focused our analysis on this significant subset of neurons, which we label “side-selective” neurons. Consistent with previous work in primate dorsal striatum (Ding and Gold, 2010), we found that the average responses of these rat striatum neurons ramped upwards for stimuli in the preferred direction (Fig. 3), and moreover, that after an initial onset latency the slope of the ramp was proportional to the stimulus strength (Fig. 3; Fig. 4a). Importantly however, a gradual ramping profile is not conclusive evidence for encoding of gradually accumulating evidence, because such a response profile can also be consistent with other encoding schemes (Ditterich, 2006; Hanks et al., 2015; Latimer et al., 2015), for example step changes in firing rate that occur at different times on different trials (Latimer et al., 2015). Thus, we extended our analysis to include a more direct test in which the influence of single quanta of sensory evidence on the responses of the cells is quantitatively assessed.

If indeed temporal integration underlies the ramping activity of the striatal cells, then each single quantum of sensory evidence (an auditory click) should result in a fixed-magnitude and sustained increase in the neuron’s firing rate (Fig. 4b, model) (Hanks et al., 2015; Huk and Shadlen, 2005). We thus estimated the effect of each sensory evidence quantum by computing the click-triggered average response of the side-selective striatal neurons. We found that striatal neurons modulated their activity in close agreement with this theoretical prediction (Fig. 4b, data), arguing in favor of a role of this anterior striatal subregion in the behavioral accumulation of evidence process.

We also took advantage of a recently developed method to compute neural tuning curves–that is, direct estimates of firing rates as a function of accumulated evidence (Hanks et al., 2015). Model-derived estimates of the moment-by-moment value of the accumulating evidence on each trial are collated with simultaneously recorded firing rates to generate tuning curves for accumulated evidence (see (Hanks et al., 2015), Methods, and illustration of the method in Extended Data Fig. 5). When applying this analysis to the striatal data we found that the side-selective neurons encoded accumulating evidence in a remarkably graded manner throughout the period of evidence accumulation (Fig. 4c, d). This graded encoding was consistent across different neurons in the population of recorded striatal cells (Fig. 5). Such a graded representation implies that the striatum carries information about the graded value of accumulated evidence, as would be required for a brain structure involved in such a process.

**Fig. 5.**
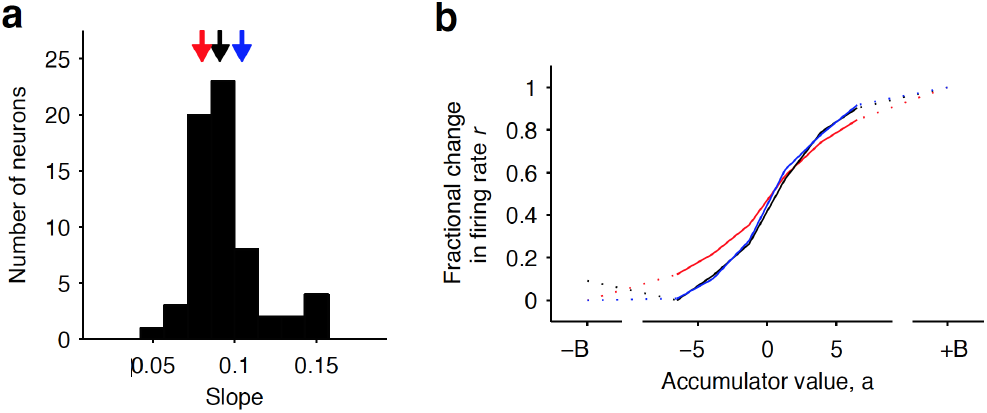
Distribution of tuning curve slopes for individual striatal neurons. **(a)** Histogram of the slope of individual neurons obtained from a sigmoidal fit of the relationship between firing rate and accumulator value. Black arrow indicates the median value of the distribution (50 percentile). Red and blue arrows indicate points corresponding to the 20 and 80 percentile mark, respectively. **(b)** Example tuning curves shown for 20, 50, and 80 percentile neurons. Graded encodings of accumulated evidence are exhibited for all of these neurons.

Our pharmacological methods address the questions of ‘Whether’ the anterior dorsal striatum is involved in the process of accumulation of evidence, and our electrophysiological and computational methods address ‘How’ the anterior dorsal striatum represents accumulating of sensory evidence. However, neither directly addresses the question of ‘When’ it is involved. This question is critical and has proven to be pivotal in assessing the involvement of a brain region in the evidence accumulation process. For example, some brain regions can be required for decisions based on accumulation of evidence, yet contribute at times that suggest they are in fact required for processes that are subsequent to the gradual accumulation itself (Erlich et al., 2015; Hanks et al., 2015). No region to date has been reported to be required at points of time that fully coincide with the evidence accumulation period.

To delineate the precise timing of the anterior dorsal striatum’s contribution, we used optogenetic inactivation, mediated by halorhodopsin eNpHR3.0, to unilaterally and transiently inactivate this region during the Poisson Clicks task. We expressed eNpHR3.0 using viral delivery methods (Fig. 6a; Methods). Acute neural recordings in our experimental rats verified that we could indeed transiently silence neural activity in the striatum at fine temporal precision using delivery of green light (Fig. 6b). We began with full-trial unilateral optogenetic inactivation and found that in agreement with the pharmacological inactivation described above, optogenetic manipulation resulted in more ipsilateral choice biases relative to control trials, which in this case were randomly interleaved with the inactivation trials (Fig. 6c; bias = 9.0 +− 2.3%, p<0.01). These effects were consistent across rats (Fig. 6d). Control rats injected into the striatum with the same virus expressing YFP alone did not show a behavioral bias (bias = 0.1 +− 1.8%, p = 0.89). Next, to directly resolve *when* the striatum contributes to the auditory accumulation of evidence task, we transiently inactivated it unilaterally during one of four different 500-ms time periods of the task: (i) the delay period immediately preceding stimulus onset (“pre-accumulation”), (ii) the first half of a 1-sec sensory stimulus (“first half”), (iii) the second half of a 1-sec sensory stimulus (“second half”), or (iv) the movement period (“post-choice”). In contrast with similar inactivation assays of the cortical Frontal Orienting Fields (FOF), which have no effect during early parts of the accumulation period(Hanks et al., 2015), we found that transient optogenetic inactivation of the anterior dorsal striatum during both the first half and second half of the accumulation caused a significant bias for the ipsilateral choices, with a similar magnitude of effect in both of these periods (first half bias = 10.4 +− 4.0, p < 0.01; second half bias = 12.9 +− 3.7%, p < 0.01; difference = 2.5 +− 2.8, p = 0.2; Fig. 6e; first half effect in striatum is significantly greater than in FOF, p < 0.01, Fig. 7). Remarkably, the effect in striatum was limited to the stimulus presentation period and we found no significant effect of optogenetic inactivation for pre-accumulation or post-choice periods (pre-accumulation bias = 0.4 +− 5.4%, p = 0.42; post-choice bias = 0.9 +− 5.2%, p = 0.38; Fig. 6e). These results are consistent with the idea that the anterior dorsal striatum plays a direct causal role throughout the entire evidence accumulation process.

**Fig. 6.**
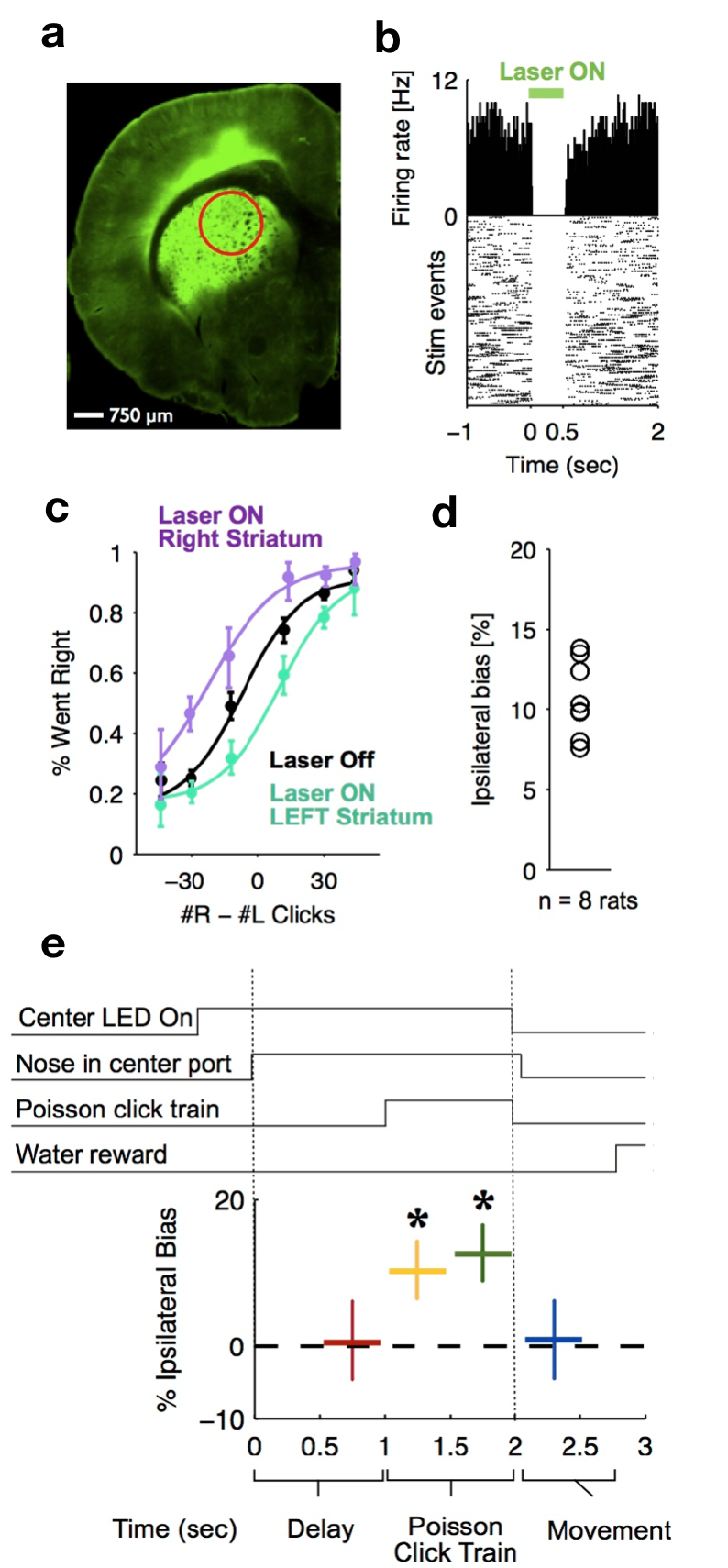
Optogenetic inactivation reveals that dorsal striatal activity causally contributes to decision formation throughout the accumulation process but not before nor after. **(a)** Coronal section of the left hemisphere showing expression of eYFP-eNpHR3.0 in the left dorsal striatum. Optical fiber localization and 750 µm estimated inactivation radius are indicated by the red circle. **(b)** Raster plot (bottom) and peri-stimulus time histogram (top) showing the effectiveness in silencing of local striatal activity in response to delivery of green light (indicated by green bar on top). **(c)** Unilateral full-trial optical inactivation of the striatum results in an ipsilateral bias on accumulation trials. Purple and cyan psychometric curves show data for right and left striatal inactivation, respectively, and black psychometric curve shows data from control trials that occurred on the same days (n = 8 rats). **(d)** Scatter plot indicated the mean ipsilateral bias for each individual rat. **(e)** Bottom: Behavioral bias caused by 500-ms inactivation during the pre-stimulus delay period (red), the first half of the sensory stimulus (yellow), the second half of the stimulus (green) and upon initiation of movement (blue). Top: Task structure. Note the significant effect (indicated by an asterisk) only during evidence accumulation but not prior to the presentation of sensory stimuli nor after.

**Fig. 7.**
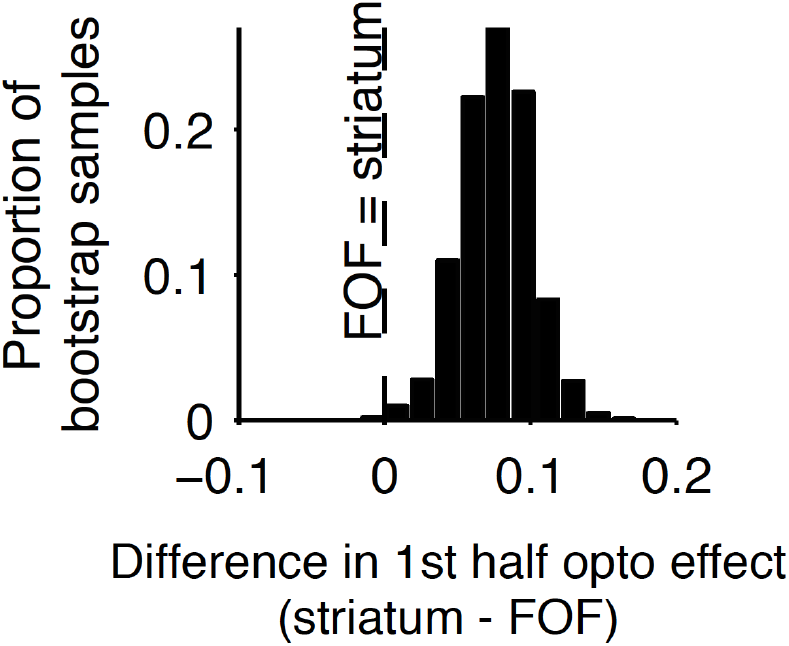
Comparison of early stimulus period optogenetic inactivation effects in striatum and frontal orienting field (FOF). Optogenetic inactivation of anterior dorsal striatum during the first half of the 1s stimulus presentation period produced a significantly larger effect than the same manipulation of the FOF (p < 0.01), with the latter data coming from a previous report. For this analysis, individual trials were resampled with replacement from both data sets across 1000 iterations, and the difference in inactivation effect was calculated for each iteration to provide a nonparametric statistical comparison. As reported above, the first-half anterior dorsal striatum effect itself is significant, and as reported previously, the first-half FOF effect is not significant, but a direct comparison as described here is still necessary to establish a significant difference.

## Discussion

Studies over more than two decades have attempted to elucidate neural circuits that underlie the accumulation of evidence over time (Carandini and Churchland, 2013; Gold and Shadlen, 2007; Krajbich et al., 2012; Shadlen and Newsome, 1996), but no brain region has previously been identified as being all of: first, necessary for accumulation-based decision-making behavior; second, having the graded neural encoding required for direct involvement in computing the graded, gradually evolving, value of the accumulating evidence; and third, making a causal contribution throughout times that fully coincide with the accumulation process. By demonstrating that the anterior dorsal striatum satisfies all three of these criteria, our work suggests the anterior dorsal striatum as the first identifiable node in the neural circuit causally responsible for computing evidence accumulation. The anterior dorsal striatum is well positioned anatomically to participate in evidence accumulation as it receives diverse convergent anatomical input from multiple cortical areas (Cheatwood et al., 2003; McGeorge and Faull, 1989) and is connected via recurrent loops with cortical and subcortical areas that are widely believed to play a role in action selection (Ding and Gold, 2013a). Whether the anterior dorsal striatum possesses a unique role in evidence accumulation, or whether it is an important node of a more extended network of brain regions that operate in coordination to mediate evidence accumulation, remains to be resolved. Corticostriatal loops are organized as distinct parallel circuits (Alexander et al., 1986; Kim and Hikosaka, 2015); future studies dissecting the contribution of different loops will be important for resolving this major question.

Our results suggest that the striatum may be directly involved in a more expansive set of computations, traditionally considered to be more cognitive in nature, than the already well-established functions of the dorsal striatum in action selection, response initiation, and habit formation (Graybiel, 2008; Hikosaka et al., 2014; Jin and Costa, 2010; Kravitz and Kreitzer, 2012). It will be important to better understand how the striatal involvement in computing accumulation of evidence, as identified in this study, may contribute to those previously established functions. The computations involved in evidence accumulation may perhaps provide an efficient mechanism for extracting important pieces of information from the environment in the service of other roles of the striatum.

Our model fits to unilateral pharmacological inactivation data found that, similar to unilateral inactivations of the FOF, the side bias in lapse rates was increased by the inactivations. But in contrast to the FOF, the accumulator became leakier and the accumulator noise was also substantially increased after striatal inactivation (Fig. 2a,b). Model fits to bilateral anterior dorsal striatum inactivation data found that the sum of sensory and accumulator noise magnitude parameters was significantly increased by striatal inactivation (Fig. 2c,d). A parsimonious account suggests that the main noise parameter affected may be the magnitude of the noise in the evidence accumulator. This is consistent with the idea that the striatum plays a role in the accumulation process. The lack of a significant effect on other parameters should be treated with caution: it remains possible that future studies with greater statistical power could discern an effect of striatal inactivation on some of these other parameters. Nevertheless, the current results do suggest accumulator noise as a principal parameter of interest. It is conceivable that bilateral striatal inactivation increases accumulator noise by destabilizing the accumulator’s representation without biasing it, but a circuit model hypothesis as to precisely how the striatum might affect the accumulator noise level remains to be developed. Another important direction for future studies will be the development of models with temporally-specific parameters that could be used to appropriately model the effects of temporally-specific optogenetic inactivation.

Independently of whether the anterior dorsal striatum operates alone or as part of a broader circuit for computing gradual evidence accumulation, and independently of precisely what its contribution to the evidence accumulation computation is, the data reported here provide a critical foothold towards delineating the relevant causal circuit: for example, the anterior dorsal striatum’s major inputs and outputs become important candidate regions to be examined for a potential role in the process. The possibility that the causal circuit for computing evidence accumulation may be delineated in the near future suggests that we will soon be able to elucidate the circuit and cellular mechanisms that support evidence accumulation, a computation that is crucial for decision-making behavior in a wide range of species, including humans.

## Supplementary Material

**This Supporting Material includes:**

Materials and Methods

### Online Methods

#### Subjects and housing

All animal procedures described in this study were approved by the Princeton University Institutional Animal Care and Use Committee and carried out in accordance with National Institutes of Health standards. All subjects were adult male Long-Evans rats (Taconic, NY) that were pair housed in Technoplast cages and were kept on 12-hr reversed light-dark cycle. All training and testing procedures were conducted during the dark cycle. Rats had free access to food but had restricted water access. The amount of water rats could obtain daily was limited to 1 hour per day of free water (starting 30-min following the end of training), in addition to what they could earn during training.

#### Behavior

Rats trained seven days a week at similar times each day for a period of about 110 minutes daily. The training took place in custom training boxes (Island Motion, NY) placed inside sound-and light-attenuated chambers (H10–25A, Coulbourn Instruments, PA). Each box consisted of three straight walls and one curved wall in which three nose ports were embedded (one in the center and one on each side, Fig. 1A). Each nose port also contained one light emitting diode (LED) that was used to deliver visual stimuli, and the front of the nose port was equipped with an infrared (IR) beam to detect the entrance of the rat’s nose into the port. A speaker was mounted above each of the side ports and was used to present auditory stimuli. Each of the side ports also contained a sipper tube that was used for water reward delivery, with the amount of water controlled by valve opening time.

All rats were trained using a semi-automated training protocol on a previously developed accumulation of evidence task(Brunton et al., 2013). Training and testing procedures were similar to those described previously(Erlich et al., 2015; Hanks et al., 2015). In brief, at the start of each trial, rats were instructed to place their nose in the central port and maintain nose fixation in response to LED illumination of that port. Subsequently, after a delay of at least 200 ms, rats were presented with a two trains of auditory clicks presented simultaneously, one from the left and one from the right speaker. For neurophysioloigcal recordings and pharmacological (muscimol) inactivation experiments, the click train duration varied between 0.1 to 1.2 s. For optogenetic experiments, the stimulus duration was fixed at 1 s for all trials. The train of clicks from each speaker was generated by an underlying Poisson process, with different mean rates for each side. The combined mean click rate was fixed at 40 Hz, and trial difficulty was manipulated by varying the click rate ratio between the two sides. The mean click rate ratio varied from 39:1 clicks/s (easiest) to 26:14 (most difficult). Upon completion of stimulus presentation, the central LED was turned off and rats had to orient towards the side that played more clicks and nose poke into the corresponding port to obtain water reward of 24 µL.

#### Surgery

The experiments described in this manuscript focus on the anterior dorsal striatum of the rat at stereotaxic coordinates of 1.9 mm anterior and 2.4 mm lateral, relative to bregma. Each rat received one of three surgical procedures that were all described in detail elsewhere for different brain areas but were identical in all other respects. These were: (i) implantation of a tetrode-based microdrive consisting of 8 tetrodes (3 rats, left striatum)(Erlich et al., 2011, 2015; Hanks et al., 2015), (ii) cannulas for pharmacological inactivation (4 rats, bilateral) (Erlich et al., 2011; Hanks et al., 2015) or (iii) chemically etched optical fibers coupled with viral injection(Hanks et al., 2015) (13 rats; 6 left striatum and 7 right striatum). The injected virus consisted of 2–3 µL of AAV virus (either AAV5-CaMKIIα-eYFP-eNpHR3.0 or AAV5-hSyn-eYFP-eNpHR3.0 or a mixture of both at a ratio of 1:2, respectively). Two of the three rats that were used for electrophysiological recordings and received a tetrode implant targeting the anterior dorsal striatum were further injected with AAV5-CaMKIIα-eYFP-ChR2 and were implanted with two optical fibers and an additional tetrode-based microdrive targeting the rat SNr, GPe and superior colliculus, respectively. This data is not discussed in the present manuscript. The infusion cannulas were implanted at an angle of 15º lateral to minimize any potential backflow of muscimol to the frontal orienting fields (FOF), which have recently been demonstrated to be necessary for maximal performance on this task (Erlich et al., 2015). Accurate placement of all implants and viral injection targeting was verified histologically.

#### Infusions

Infusion procedures follow methods described in detail previously (Erlich et al., 2015). Briefly, infusions were generally performed during normal training sessions, were usually at least one week apart, and never on consecutive days. Control sessions took place on the day prior to the infusion session. On the day of infusion, rats were lightly anesthetized with 2% isoflurane and anesthesia was sustained via continuous delivery of isoflurane using a nose cone. Using a Hamilton syringe that was attached via tubing to the injector, we delivered 0.5 µL of muscimol at concentration of 0.125 mg/mL to either the left or right side of the anteriodorsal part of the rat striatum during unilateral infusion sessions and to both sides during bilateral infusion sessions (Fig. 1 B and C, respectively). After delivery, the injector was left inside the brain for a minimum period of 5 minutes to allow adequate diffusion before removal and also to minimize backflow along the cannula tract. Subsequently, the injector was removed, the cannula was closed, and the rat was removed from isoflurane and placed back into its home cage. We allowed 30 minutes of recovery from anesthesia before placing the rat into the behavioral box.

#### Optogenetic perturbation

The methods used in this study for optogenetic perturbation are identical to those described in detail previously (Hanks et al., 2015). Prior to each experimental session, a 532 nm green laser (OEM Laser Systems) was connected via a 1m patch cable with a rotary joint (Princetel) and FC connector to the rat’s optical implant. The rotary joint was mounted on the ceiling of the behavioral chamber. The laser operated at 25 mW and was triggered by a 5V transistor-transistor logic (TTL) pulse, delivered in response to behavioral events and triggered by the automated traiwhayning software (BControl). On all experimental days, laser illumination occurred on a random subset (25%) of trials and applied unilaterally. Laser illumination trials could be divided into two main types. In the first, we delivered light for a continuous period of 2s, starting 500 ms prior to the initiation of the auditory clicks stimulus and ending 500 ms after the termination of the click train. This trial type is defined as ‘full-trial’ inactivation. For this we used a cohort of 8 rats. In the second trial type, which we refer to as ‘time-resolved’ inactivation, light illumination was delivered in one of four different 500 ms time periods: the delay before stimulus onset, the first half of the 1 s auditory stimulus, the second half of the 1 s auditory stimulus, or during the movement period (Fig. 3E). All time resolved inactivation periods were randomly interleaved within single behavioral sessions. For ‘time-resolved’ inactivation experiments, we used a cohort of 7 rats, 2 of which also belonged to the full-trial inactivation cohort.

The physiological effect of eNpHR3.0 on local neuronal activity was tested using acute recordings in experimental rats (Fig. 3A), as described previously(Hanks et al., 2015). Rats were anesthetized using isoflurane and a sharp etched optical fiber was inserted into the center of the field of viral infection. The optical fiber was coupled with a 532 nm green laser with ~25 mW light intensity at the tip. In parallel, a sharp tungsten electrode (1 MΩ) was positioned adjacent to the optical fiber tip. The effect of laser activation on spontaneous activity was tested by The physiological effect of eNpHR3.0 on local neuronal activity was tested using acute delivering a series of pulses, 500 ms duration each, at 25 mW every 5 s. The signals from the electrode were amplified, filtered (300–6000 Hz), thresholded based on voltage (30 µV) and sampled at 30.3 kHz (0.25 ms before the threshold triggering and 0.75 ms after; Neuralynx Cheetah). The spikes and TTL pulses were time-stamped with the same 1-MHz clock (Digital I/O, Neurlaynx).

#### Neural recording and spike sorting

Neural recordings and spike sorting methods were previously described in detail(Erlich et al., 2011; Hanks et al., 2015). Briefly, over the course of ~2–4 weeks following surgery, the tetrodes were slowly lowered towards the dorsal part of the rat anterior striatum. On most recording days, an electrically-quiet electrode was used as a reference channel, and in the cases that such an electrode was not available, we used the ground of our neurophysiology recording system (Nerualynx) for reference. During recording, a unity-gain preamplifier (HS-32, Neuralynx) was attached to a connector on top of the microdrive via a light-weight tether. Signals from each of the channels were amplified (1,400–5,000×) and band-pass filtered (300–6,000 Hz; Digital-Lynx, Neuralynx). A voltage threshold (20–50 µV) was used for collecting 1-ms spike waveforms, which were sampled at 30.3 kHz (0.25 ms before the triggered event and 0.75 ms after; Neuralynx Cheetah). Neural activity was recorded daily during behavioral sessions that lasted 2–4 hours on average. Regardless of the quality of the recordings, tetrodes were never kept in the same position between days, and were always moved at the end of each recording day (40– 200 µm daily), in order to obtain recordings from new ensembles of neurons daily.

#### Analysis of causal perturbation data – Optogenetics and pharmacological inactivation

Detailed methods for generating psychometric curves and estimating biases resulting from inactivation in rats performing this exact behavioral task were recently described(Erlich et al., 2015; Hanks et al., 2015).

In brief, for muscimol inactivation experiments, psychometric curves were generated by concatenating data across either infusion or control sessions for individual rats and fitting a 4-parameter sigmoid described using the following formula: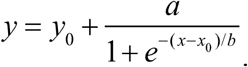.

In that equation, the ‘x’ variable is the click difference on each trial (# Right Clicks – # Left Clicks), ‘y’ is the proportion of trials on which the animal went ‘Right’, and the four parameters of the fitting procedure are: ‘x_0_’, the inflection point of the sigmoid; ‘b’, the slope of the sigmoid; ‘x_0_’ and ‘a + y_0_’, the minimum and maximum of the proportion on which the rat went ‘Right’, respectively.

For optogenetic inactivation experiments, we measured behavioral bias resulting from transient inactivation of neural activity on a subset of trials (25%) by first binning the trials based on stimulus strength. We then computed the mean difference between the fraction of trials that the rats went to the side ipsilateral to side of its optical implant for inactivation and control trials for each of 10 binned stimulus strengths. Thus, a positive value resulting from this measurement represents an increase in ipsilateral responses on laser illumination trials over control trials where optical stimulation was absent. Confidence intervals and statistical comparisons for this bias parameter were calculated using nonparametric bootstrap procedures. The bias resulting from unilateral pharmacological inactivation was calculated in a similar way, but the control behavior was derived from non-inactivation control sessions obtained the day before inactivation. The performance impairment resulting from bilateral pharmacological inactivation sessions was also calculated using the non-inactivation control sessions obtained the day before inactivation. Performance was defined as percent correct trials for each binned stimulus strength.

#### Analysis of neural recording

Spike waveforms were sorted on the basis of their relative energies and amplitudes on different channels of each tetrode. Clustering software (SpikeSort3D, Neuralynx) was used to manually isolate single units. Each spike was graphically positioned in a two-or three-dimensional space representing the energy or amplitude of the spike on two or three of the four tetrode channels. Convex hull boundaries and template-matching of waveforms were used to identify well-separated clusters of spikes, which were individually color coded. Data from the entire session were spike-sorted together. To compute the peri-event time histogram (PETH) for the population activity in response to the presentation of auditory clicks (Fig. 2A & Extended Data Fig. 4) we followed the following procedure. For all well-isolated single units, individual trial rate functions were first generated by smoothing the spike trains with a causal half-Gaussian filter with 0.1 s standard deviation. The response functions of individual neurons were then normalized based on the mean firing of each individual neuron at the time of stimulus onset. Trials were subsequently sorted by a quantity that we defined as the ‘mean stimulus strength’ following the same procedure that has been described previously(Hanks et al., 2015). Mean stimulus strength was defined by dividing trials for each neuron into quantiles based on difference of the preferred and non-preferred click rates.

The influence of single auditory clicks on neural responses, the ‘click-triggered average’, was calculated as followed. The trials of individual neurons were first grouped based on the underlying Poisson rates that were used to generate the auditory stimuli. For each group, the mean PETH was computed. This quantity corresponds to the expected neural response at each point in time for each Poisson rate group. This mean response was then subtracted from each trial to generate the residual response from the expected one given the Poisson rate. Aligning this residual response to a click describes the change in the neural response that is associated with a single auditory click relative to the average expected response to clicks at other times. These click-aligned residual responses were averaged across all click times to obtain the click-triggered average response for each Poisson rate group. The click-triggered average for each neuron was calculated by averaging across the different Poisson rate groups. To compute the response across clicks arriving from the both preferred and non-preferred sides, we inverted the residual response for non-preferred direction clicks prior to averaging. The click-triggered average response profiles generated using this procedure were compared to a model-based prediction based on a graded, linear encoding of accumulated evidence. To do this, we simulated evidence accumulation trajectories for 5,000 trials using the same range of stimulus difficulties and durations that existed for the neural data. We then encoded these simulated trajectories with a graded, linear function of accumulated evidence (firing rate *r* = *k_1_ x a(t)* + *k_2_* in which k_1_ and k_2_ are constants). Finally, we applied the same analysis described for the neural data to estimate the predicted click-triggered average under this encoding (Fig. 2B).

#### Behavioral model-based analysis of neural data

We applied recently developed methods in our lab to generate tuning curves that specify the relationship between neural firing rates and mentally accumulated evidence(Hanks et al., 2015). These techniques take advantage of a behavioral model that provides a moment-by-moment and trial-by-trial estimation of the mentally accumulated evidence for this task (Brunton et al., 2013; Hanks et al., 2015). The model converts each trial’s incoming stream of discrete left and right clicks into an accumulating evidence quantity *a(t)* that determines choice behavior. Parameters that govern how *a(t)* evolves are fit based on the rat’s behavior, with the choices made on individual trials constraining the possible trajectories of *a(t)*. Thus, on each trial, the model estimates the evolution of a noise-induced probability distribution over values of the accumulating evidence *a(t)*.

A full description of the model (Brunton et al., 2013; Erlich et al., 2015) is as follows:

At each time point, the accumulator memory *a(t)* represents an estimate of the right versus left evidence accrued so far. At stimulus end, the model decides right if *a* > Ϸ and left otherwise, where Ϸ is a free parameter. Right (left) pulses change the value of *a* by positive (negative) impulses of magnitude *C*. σ_a_^2^ is a diffusion constant, parameterizing noise in a. σ_s_^2^ parameterizes noise when adding the evidence from a right or left pulse: For each click, variance σ_s_^2^ is scaled by the amplitude of *C* and then added to the evidence contributed by the click. λ parameterizes consistent drift in the memory *a*. In the “leaky” or forgetful case (λ < 0), drift is toward *a* = 0, and later pulses affect the decision more than earlier pulses. In the “unstable” or impulsive case (λ > 0), drift is away from *a* = 0, and earlier pulses affect the decision more than later pulses. The memory’s time constant τ = 1/λ. B is the height of the sticky decision bounds and parameterizes the amount of evidence necessary to commit to a decision. ϕ and τ_ϕ_ parameterize sensory adaptation by defining the dynamics of *C*. Immediately after a click, the magnitude *C* ismultiplied by ϕ. *C* then recovers toward an unadapted value of 1 with time constant τ_ϕ_. Facilitation is thus represented by ϕ > 1, whereas depression is represented by ϕ < 1.

These properties are implemented by the following equations if |*a*| ≥ B then 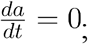 else 
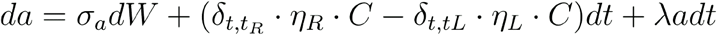
where δ_*t,t*_*R,L*__ are delta functions at the times of the pulses; *η* are i.i.d. Gaussian variables drawn from *N*(1,σ_*s*_ ; and *dW* is a white-noise Wiener process. The initial condition *a*(*t* = 0) is 0.

Adaptation dynamics are given by 
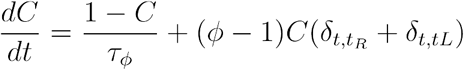

In addition, a lapse rate parameterizes the fraction of trials on which a random response is made. Ideal performance (*a* = #right clicks − #left clicks) would be achieved by 
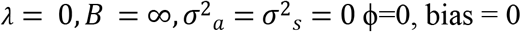

This estimate *a(t)* was then related directly to neural firing rates on individual trials to estimate neural tuning curves for accumulating evidence. The estimates of the neural response and accumulating evidence on individual trials were used to calculate the joint probability distribution between those two variables as a function of time for each neuron. The correspondence between time in the model and neural time was determined based on the latency of the stimulus-dependent response modulation. This latency was calculated as the first time bin the PETH to have a significant modulation of neural response based on stimulus strength, which corresponded to 150 ms (Fig. 2A). Thus, t = 0 in the model was taken as 150 ms after stimulus onset. From the joint probability, we extracted each neuron’s response conditional on the value of the accumulator. We then combined across neurons by weighting the contribution of each by the inverse of the variance of this conditional distribution, which gives more weight to representations that are less noisy.

To quantify the relationship between neural response and accumulator value across time, we averaged across the time period from 0.15 to 0.5 s into the decision process. To characterize the encoding across individual neurons, we fit this relationship of the response to the accumulator value with a four-parameter sigmoid using the following equation: 
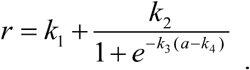

In this equation, k_2_k_3_/4 determines the slope at zero-crossing, which characterizes whether the neural response changes smoothly between negative and positive accumulator values or whether it changes sharply in this region.

#### Histology

The rat was fully anesthetized with 0.4 mL ketamine (100mg/ml) and 0.2 mL xylazine (100mg/ml) IP, followed by transcardial perfusion of 100 mL saline (0.9% NaCl, 0.3 × PBS pH 7.0, 0.05 mL heparin 10,000 USP units/mL), and finally transcardial perfusion of 250 mL 10% formalin neutral buffered solution (Sigma HT501128). The brain was removed and post fixed in 10% formalin solution for a minimum period of 7 days. 100 micrometer sections were prepared on a Leica VT1200S vibratome, mounted on Superfrost Pus glass slides (Fisher) with Fluoromount-G (Southern Biotech) mounting solution and glass cover slips. Images were acquired on a Nikon Eclipse Ti fluorescence microscope under 4x magnification.

#### Four model parameters to quantify sources of a lateralized bias

The original model had only a single parameter able to describe a right versus left choice bias, the decision borderline Þ. By adding three more parameters that could cause different types of side biases, fitting the extended model to behavioral data following a unilateral inactivation, and asking which parameters are most affected relative to control trials, we can estimate which particular aspect of the behavior was impacted by the inactivation. The 4 different sources of a choice bias that we considered were:

**Post-categorization bias:** Accumulator value is categorized into ‘Go Left’ or ‘Go Right’ decisions according to a > Þ versus a < Þ, where Þ is the decision borderline parameter. When performing unilateral inactivation, the choice directions can be mapped as “contralateral” or “ipsilateral” with respect to the side of inactivation. Contralateral lapse rate is a fraction of the trials categorized as choices contralateral to the inactivated side of the brain, and converts them into ipsilateral choices. And ipsilateral lapse rate is a fraction of the trials categorized as choices ipsilateral to the inactivated side of the brain, and converts them into contralateral choices. The scaling is biased when κ_C_ ≠ κ_I_. So, we re-parametrize lapse rate parameters for each side as a total lapse (κ_C_ + κ_I_) and a biased lapse (κ_C_ – κ_I_).

**Input gain bias:** This can be thought of as a form of sensory neglect: Left and Right clicks have different impact magnitudes on the value of the accumulator. 0.5 is the balanced point, where left and right clicks have same impact magnitudes. If the value of input gain weight is lower than 0.5, then ipsilateral clicks have a stronger impact, and decision will consequently be biased to the ipsilateral side. The closer the value to 0, the impact of ipsilateral clicks gets stronger. While, the closer the value to 1, the impact of contralateral clicks gets stronger.

**Accumulation shift:** Before categorizing the accumulator into ‘Go Left’ vs ‘Go Right’ decisions (by comparing the accumulator’s value to 0), a constant is added to the value of the accumulator. In the behavioral model, this is implemented by changing a to a + shift after the end of accumulation but before the application of the decision borderline, or (equivalently) by changing the decision borderline Þ to Þ – shift, with shift being the free parameter, fit to the behavioral data.

**Biased sensory noise:** By differentially affecting signal-to-noise rations from the two sides, can be thought of as a form of sensory neglect distinct from input gain bias: Left and Right clicks have different magnitudes of noise in their impact. The biased sensory noise was implemented by allowing the contralateral noise variance to be a free parameter, fit to the behavioral data from unilateral inactivation trials.

**Extended Data Fig. 1.**
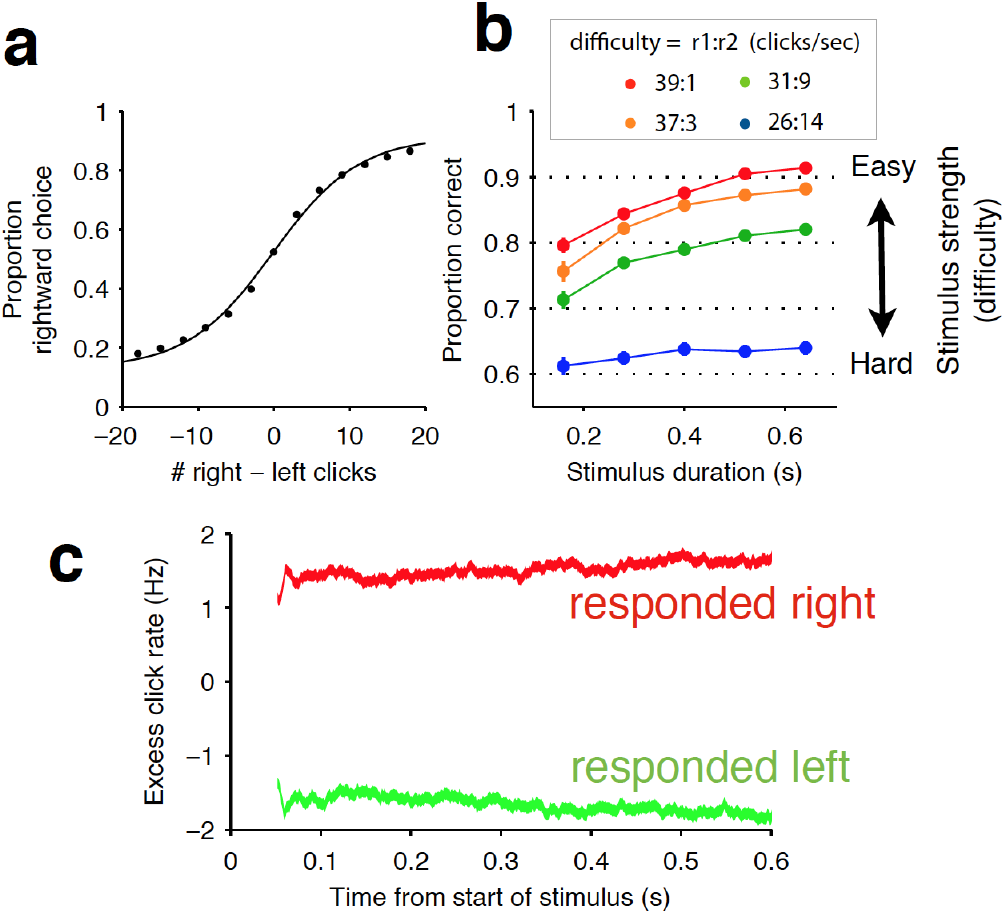
Rat behavior indicates accumulation of auditory evidence over the entire trial. **(a)** Averaged psychometric curve showing average performance across all rats during the Poisson-clicks task (n = 20 rats). **(b)** Chronometric curve of the same rats showing improved performance with longer stimulus durations. Difficulty is grouped by the ratio of auditory clicks played on the left and right sides. **(c)** Psychophysical reverse correlation analysis. Red and green curves correspond to trials on which rats went right and left, respectively. Note that stability of each curve indicating that sensory evidence (auditory clicks) throughout the trial were weighted evenly in the rats’ decision process. This result is further corroborated with the long accumulation time constants exhibited by the rats (see Extended Data Table 1).

**Extended Data Fig. 2.**
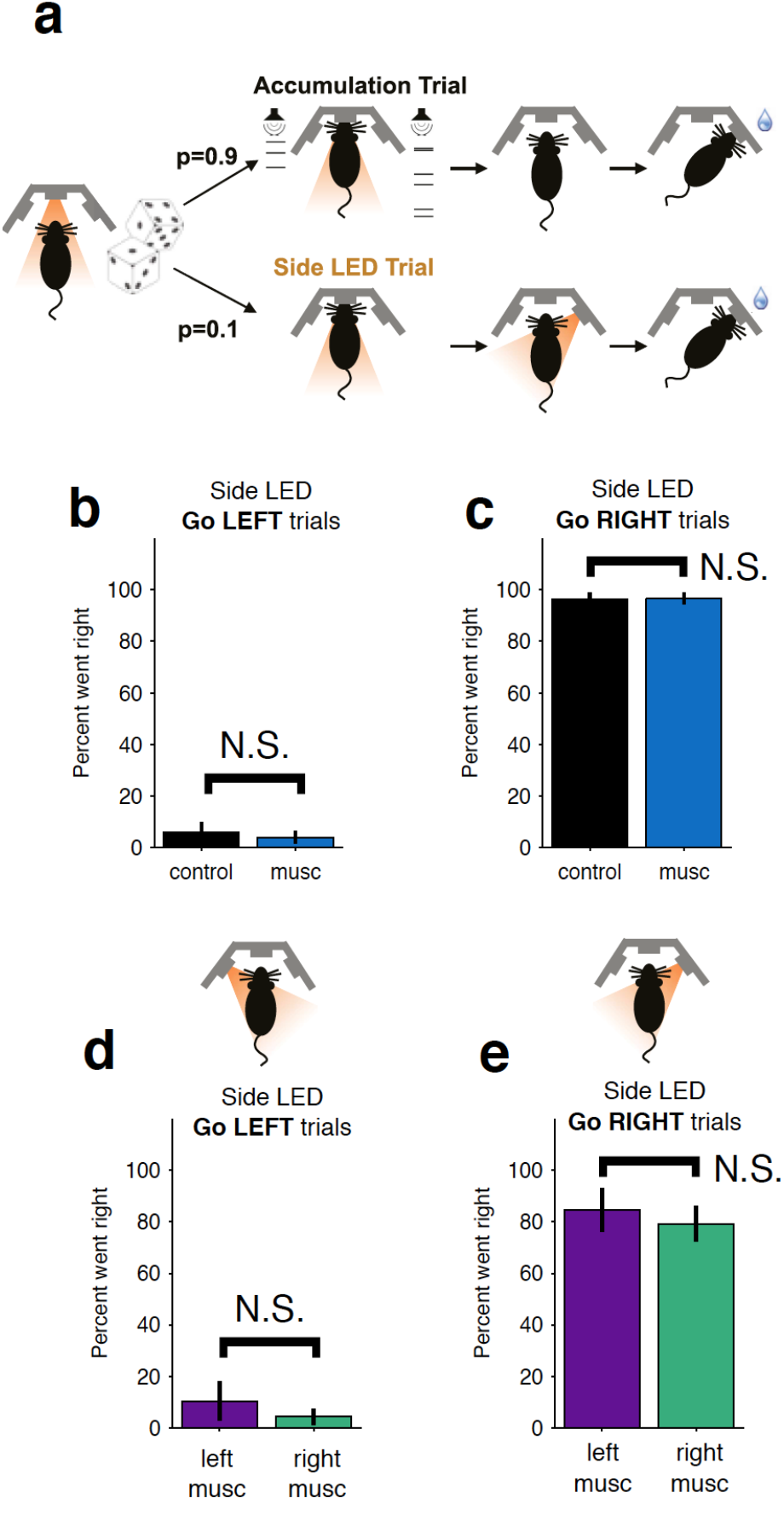
Control LED trials indicate the behavioral bias and impairments resulting from striatal inactivation are not due to motor impairments. **(a)** On ~10% of trials, rats were presented with an LED above the right or left port and had to orient towards that port. Thus, they had to perform the same motor action but success was not dependent on evidence accumulation. No significant impairment is observed for either bilateral inactivation **(b-c)** or unilateral **(d-e)** inactivations. Color scheme is the same as in Fig. 1 b-c.

**Extended Data Fig. 3.**
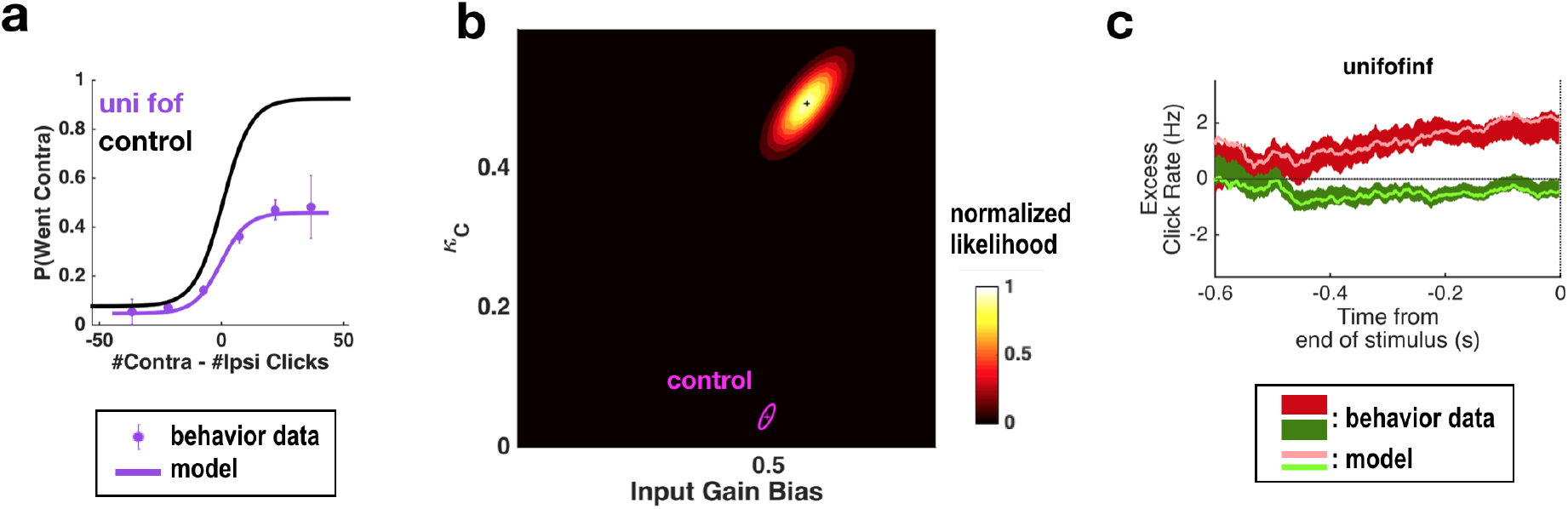
Fitting all parameters simultaneously for unilateral FOF inactivation data confirms the same conclusions as previously found fitting only two parameters at a time. **(a)** Psychometric curves for control and unilateral FOF inactivation data. The black line is the model fit to the control data. The purple circles with error bars are experimental data from unilateral FOF inactivation sessions, and indicate fraction of Contra choice trials (mean ± binomial 95% conf. int.) across trial groups, with different groups having different #Contra − #Ipsi clicks. The purple line is the psychometric curve generated by the post-categorization bias model. **(b)** The 2-dimensional normalized likelihood surface. The peak of the likelihood surface for the inactivation data is significantly different from control for post-categorization bias (from 0.044 to 0.4933). Input gain bias (which is parametrized so that 0.5 means no side bias) ris not significantly different from its control value. **(c)** Reverse correlation analyses showing the relative contributions of clicks throughout the stimulus in the rats’ decision process. The thick dark red and green lines are the means ± std. err. across trials for contralateral and ipsilateral trials. Thin light red and green lines are the reverse correlation traces generated by extended model.

**Extended Data Fig. 4.**
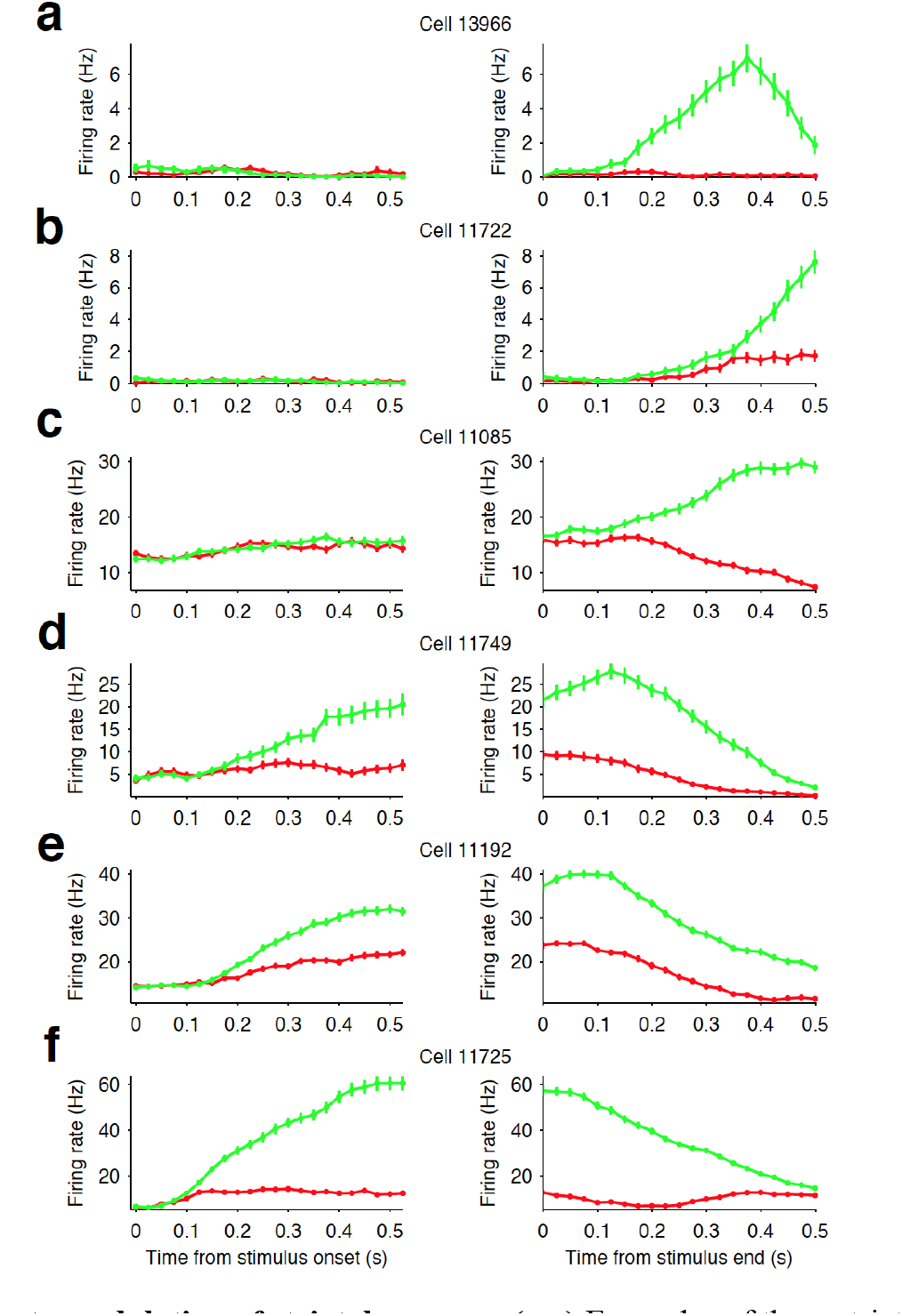
Firing rate modulation of striatal neurons. **(a-c)** Examples of three striatal neurons that did not exhibit significant modulation in their firing rates during stimulus presentation (Left column) but did show movement-related firing rate modulation (right column). **(d-e)** Examples of three striatal neurons that exhibited modulation of their firing rate during stimulus presentation and exhibited side-selective responses. This later class of neurons are the subject of our analyses in this manuscript.

**Extended Data Fig. 5.**
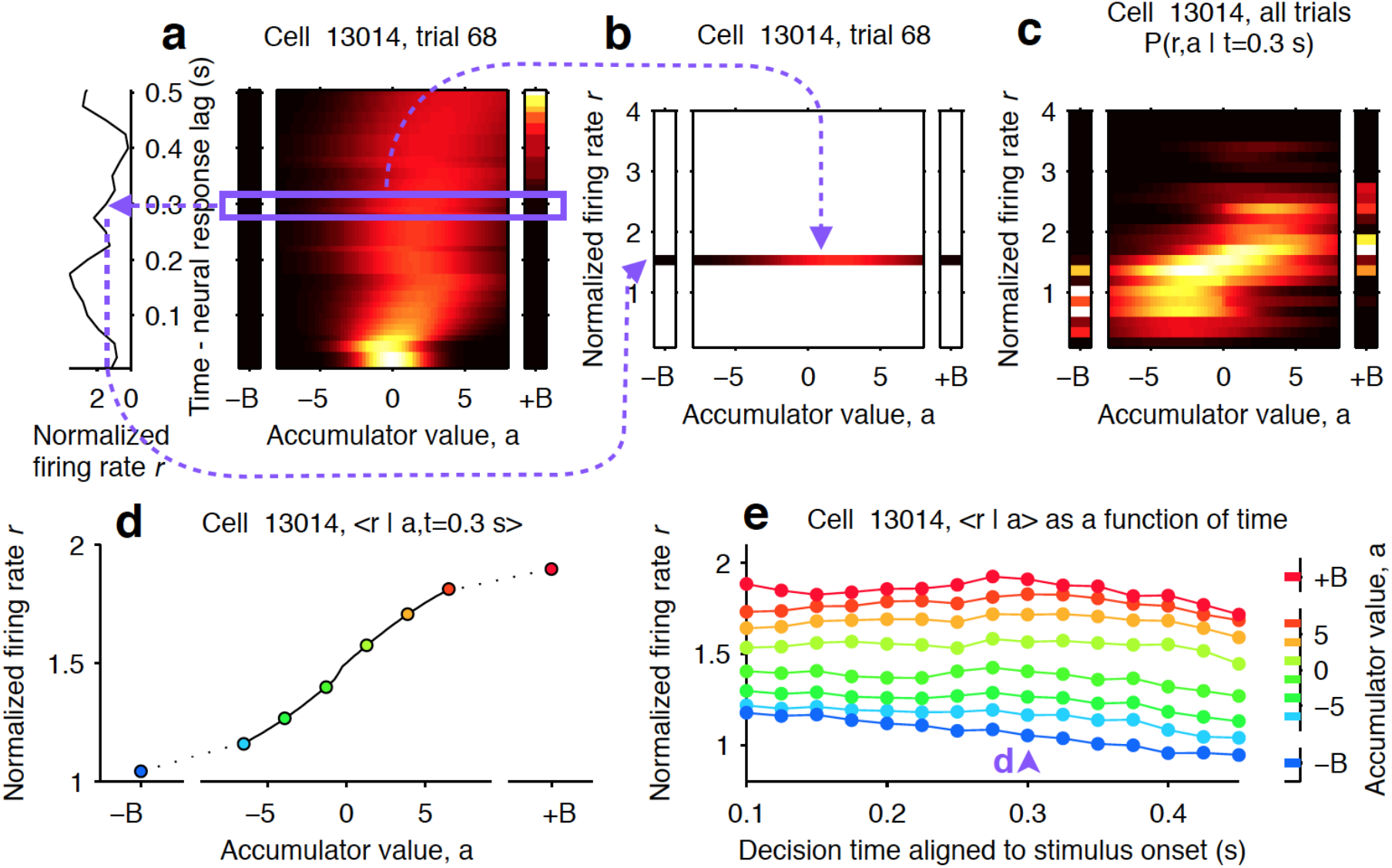
Computing tuning curves that describe the relationship between neural activity and accumulated evidence. **(a)** One trial for an example neuron from the striatum. The left side shows the firing rate of the neuron, and the right side shows the behavioral model’s estimate of the evolution of the distribution of the accumulator value, a (color represents probability density). Time runs vertically and is aligned to stimulus onset minus neural response lag (see methods). ±B correspond to the ‘sticky’ decision-commitment bounds on evidence accumulation. **(b)** Building a map of firing rate versus accumulator value. At a given time point (here, t = 0.3 s), we copy the distribution of a (blue box) to a vertical position given by the firing rate of the neuron. **(c)** Continuing with the same time point, we add a slice from every recorded trial. This produces the full joint distribution P(r,a | t = 0.3), the probability of seeing firing rate r and accumulator value a at time t = 0.3 s. **(d)** The accumulator values are binned, and the mean firing rate is computed for each bin to generate a neural tuning curve as a function of the accumulator value a. **(e)** The process is repeated for each time point. Each vertical slice corresponds to a tuning curve, with the one from d shown above the blue arrowhead.

**Extended Data Table 1.** Best fit parameters of behavioral model.

| Experiment | Data set | $\lambda$ | $\sigma_a^2$ | $\sigma_s^2$ | $B$ | $\phi$ | $\tau_\phi$ | p | lapse |
| --- | --- | --- | --- | --- | --- | --- | --- | --- | --- |
| pharmacological inactivation | E005 | -1.30 | 0.5 | 30.0 | 15.2 | 0.01 | 0.21 | 0.29 | 0.10 |
| pharmacological inactivation | T089 | -1.47 | 6.6 | 152.7 | 16.4 | 0.01 | 0.07 | -0.57 | 0.05 |
| pharmacological inactivation | T097 | -1.71 | 3.5 | 278.5 | 17.4 | 0.50 | 0.09 | 0.00 | 0.37 |
| pharmacological inactivation | T098 | -1.87 | 48.4 | 45.5 | 8.6 | 0.01 | 0.06 | 0.63 | 0.00 |
| electrophysiology | E021 | -0.15 | 0.0 | 40.2 | 16.4 | 0.13 | 0.04 | -0.01 | 0.10 |
| electrophysiology | T080 | 0.74 | 20.9 | 15.2 | 17.7 | 0.81 | 0.50 | -0.01 | 0.10 |
| electrophysiology | T103 | 0.30 | 73.4 | 200.0 | 15.3 | 0.10 | 0.01 | 0.86 | 0.11 |
| electrophysiology | physiology combined rat | -1.49 | 0.0 | 200.0 | 8.5 | 0.28 | 0.01 | 0.51 | 0.08 |
| optogenetics | E006 | 0.87 | 7.2 | 15.8 | 5.8 | 0.52 | 0.12 | 0.80 | 0.17 |
| optogenetics | E007 | -1.64 | 4.1 | 60.2 | 6.5 | 0.33 | 0.04 | 1.20 | 0.18 |
| optogenetics | E008 | -0.80 | 0.0 | 94.6 | 24.8 | 0.55 | 0.05 | -1.65 | 0.23 |
| optogenetics | E044 | -1.50 | 5.4 | 29.2 | 7.7 | 0.84 | 0.50 | -2.37 | 0.08 |
| optogenetics | E048 | -0.67 | 0.1 | 149.1 | 31.8 | 0.94 | 0.50 | -1.32 | 0.02 |
| optogenetics | E049 | 0.11 | 13.1 | 21.9 | 31.1 | 0.81 | 0.32 | -1.40 | 0.11 |
| optogenetics | E054 | -1.55 | 26.4 | 183.6 | 26.2 | 0.94 | 0.50 | -0.10 | 0.00 |
| optogenetics | E064 | -0.48 | 30.0 | 133.0 | 31.2 | 0.92 | 0.50 | -1.29 | 0.03 |
| optogenetics | E065 | -1.44 | 48.5 | 193.3 | 32.0 | 0.94 | 0.50 | -4.13 | 0.03 |
| optogenetics | E067 | -1.24 | 0.0 | 94.6 | 23.8 | 0.93 | 0.50 | -0.70 | 0.03 |
| optogenetics | E080 | -0.85 | 8.5 | 41.6 | 4.5 | 0.30 | 0.03 | 1.13 | 0.13 |
| optogenetics | J194 | -1.18 | 8.2 | 199.9 | 32.0 | 0.95 | 0.50 | 2.21 | 0.07 |
| optogenetics | T107 | -1.31 | 4.2 | 200.0 | 32.0 | 0.93 | 0.50 | -0.87 | 0.00 |

**Extended Data Table 2.** Best fit parameters (unilateral striatum inactivation data)

| | $\lambda$ | $\sigma^2_a$ | $\sigma^2_{s,l}$ | $\sigma^2_{s,c}$ | **B** | $\phi$ | $\tau_\phi$ | **p** | $\kappa_c + \kappa_l$ | $\kappa_c - \kappa_l$ | input gain |
| --- | --- | --- | --- | --- | --- | --- | --- | --- | --- | --- | --- |
| uni-striatum control | -1.18 | 0 | 34.68 | 53.36 | 12.66 | 0.18 | 0.29 | -0.02 | 0.29 | 0 | 0.47 |
| uni-striatum | -3.62* | 115.44* | 81.58 | 4.55 | 8.57 | 0.01 | 0.06 | 1.17* | 0.14 | 0.14* | 0.37 |

**Extended Data Table 3.** Best fit parameters (bilateral striatum inactivation data)

| | $\lambda$ | $\sigma^2_a$ | $\sigma^2_s$ | **B** | $\phi$ | $\tau_\phi$ | **p** | $\kappa$ |
| --- | --- | --- | --- | --- | --- | --- | --- | --- |
| bi-striatum control | -0.43 | 0.002 | 7.37 | 6.16 | 0.54 | 0.75 | -0.07 | 0.37 |
| bi-striatum | 0.06 | 85.66* | 21.06* | 13.25 | 0.01 | 0.04 | -0.56* | 0.51 |

**Extended Data Table 4.** Best fit parameters (unilateral FOF inactivation data)

| | $\lambda$ | $\sigma^2_a$ | $\sigma^2_{s,l}$ | $\sigma^2_{s,c}$ | **B** | $\phi$ | $\tau_\phi$ | **p** | $\kappa_c + \kappa_l$ | $\kappa_c - \kappa_l$ | input gain |
| --- | --- | --- | --- | --- | --- | --- | --- | --- | --- | --- | --- |
| uniFOF control | 1.25 | 0 | 50.53 | 70.29 | 7.87 | 0.23 | 0.11 | 0.03 | 0.09 | -0.01 | 0.50 |
| uniFOF | -0.85 | 0.93 | 71.78 | 87.10 | 11.47 | 0.27 | 0.09 | 0.29* | 0.52* | 0.46* | 0.55 |

**Extended Data Table 5.** Best fit parameters (bilateral FOF inactivation data)

| | $\lambda$ | $\sigma^2_a$ | $\sigma^2_s$ | **B** | $\phi$ | $\tau_\phi$ | **p** | $\kappa$ |
| --- | --- | --- | --- | --- | --- | --- | --- | --- |
| BiFOF control | 1.87 | 0.02 | 0.0001 | 14.40 | 0.34 | 0.75 | 0.04 | 0.17 |
| BiFOF | -4.08* | 63.18 | 151.17 | 15.74 | 0.08 | 0.04 | 0.72* | 0.09 |

